# Accurate Assembly of Circular RNAs with TERRACE

**DOI:** 10.1101/2024.02.09.579380

**Authors:** Tasfia Zahin, Qian Shi, Xiaofei Carl Zang, Mingfu Shao

**Affiliations:** Department of Computer Science and Engineering, The Pennsylvania State University, University Park, PA 16802, USA; Huck Institutes of the Life Sciences, The Pennsylvania State University, University Park, PA 16802, USA

**Keywords:** Circular RNA, Assembly, RNA-seq

## Abstract

Circular RNA (circRNA) is a class of RNA molecules that forms a closed loop with its 5’ and 3’ ends covalently bonded. Due to this specific structure circRNAs are more stable than linear RNAs, admit distinct biological properties and functions, and have been proven to be promising biomarkers. Circular RNAs were severely overlooked previously owing to the biases in the RNA-seq protocols and in the detection algorithms, but recently gained tremendous attentions in both aspects. However, most existing methods for assembling circRNAs heavily rely on the annotated transcriptomes, and hence exhibit unsatisfactory accuracy when a high-quality transcriptome is unavailable. Here we present TERRACE, a new algorithm for full-length assembly of circRNAs from paired-end total RNA-seq data. TERRACE uses the splice graph as the underlying data structure to organize the splicing and coverage information. We transform the problem of assembling circRNAs into finding two paths that “bridge” the three fragments in the splice graph induced by back-spliced reads. To solve this formulation, we adopted a definition for optimal bridging paths and a dynamic programming algorithm to calculate such paths, an approach that was proven useful for assembling linear RNAs. TERRACE features an efficient algorithm to detect back-spliced reads that are missed by RNA-seq aligners, contributing to its much improved sensitivity. It also incorporates a new machine-learning approach that is trained to assign a confidence score to each assembled circRNA, which is shown superior to using abundance for scoring. TERRACE is compared with leading circRNA detection methods on both simulations and biological datasets. Our method consistently outperforms by a large margin in sensitivity while maintaining better or comparable precision. In particular, when the annotations are not provided, TERRACE can assemble 123%-412% more correct circRNAs than state-of-the-art methods on human tissues. TERRACE presents a major leap on assembling full-length circRNAs from RNA-seq data, and we expect it to be widely used in the downstream research on circRNAs.

## 1 Introduction

Splicing is a ubiquitous and essential post-transcriptional modification of precursor mRNAs. In this process, introns are excised and the two flanking exons are stitched together. Canonical splicing covalently connects the 3’ end of an upstream exon to the 5’ end of the immediate downstream exon. A class of noncanonical splicing, known as back-splicing, stitches the 3’ end of a downstream exon to the 5’ end of an upstream exon via a back-splicing junction (BSJ), forming a closed circular structure called circular RNA (circRNA). Both canonical and back-splicing junctions may experience alternative splicing; consequently, a gene can express multiple circRNAs [10,16]. CircRNAs are more prevalent and conserved than previously thought. More than 60% of human genes express at least one circRNAs [10]. Notably, BIRC6, a gene that has only 12 linear transcripts, was found to express 243 circRNAs [10]. CircRNAs usually have a lower expression level than their linear forms, but certain circRNAs are significantly more abundant and constitute the major isoform of their genes [27,10,24,13]. Expression of circRNAs is also often tissue and developmental stage-specific in a spatiotemporal pattern [25,21,19].

An increasing volume of research has been evidencing the regulatory functionality of circRNAs and their use in disease diagnosis [13,19]. To name a few, circBIRC6 has been found to play a role in cell pluripotency [32], similarly, circHIPK3 in cell growth [37] and circular Foxo3 in suppression of cancer cell proliferation and survival [31]. Due to its circular structure, circRNAs are resistant to most RNA degradation mechanisms [16].

In particular, its lack of free 5’ and 3’ ends protects the molecule itself from exonucleases [23], making them more stable and have a longer half-life than their linear counterparts [9,8]. CircRNAs have been reported to serve as novel biomarkers in carcinogenesis and pathogenesis [21,13,28] and have found their potential use in non-invasive diagnosis [26].

It is therefore of great interest to detect expressed circRNAs in cells. The RNA sequencing (RNA-seq) protocols that target on linear RNAs or mRNAs with poly-A tails will neglect circRNAs [13]. Tailored RNA-seq experiments have been designed for circRNAs, for example, using RNase R to digest linear RNAs and therefore to enrich circRNAs followed by sequencing [10,26]. These approaches are efficient in raising the sensitivity of circRNA detection, but often low-throughput and costly. The total RNA-seq technology, on the other hand, can capture and sequence the entire population of RNA molecules, including circRNAs, in a biological sample. Total RNA-seq is high-throughput, and has been widely used in many studies about RNAs. Large-scale total RNA-seq datasets are available in public repositories such as GEO [2], SRA [14], and ENCODE [3], providing a rich resource to further study circRNAs.

Numerous computational methods to detect circRNAs from total RNA-seq were published lately (see [27] for a recent review). However, many of them require a fully annotated transcriptome, including CYCLeR [22], psirc [33], CircAST [29], CIRCexplorer2 [36], and CIRCexplorer3 [18]. Those methods’ dependency on an existing annotation significantly limits their capability to detect novel circRNAs and their applicability to non-model species without a well-annotated transcriptome. Other tools, including CIRI-full [38], Circall [20], CircMiner [1], CIRI2 [7], CIRI-AS [6], and CircMarker [17], can be operated annotation-free, but the functionality of many of them (except CIRI-full as far as we know) are constrained to identifying BSJ only, in a deficiency of assembling full-length circRNAs. Additionally, some methods have to be combined with experimental enrichment of circRNA [22]. Due to the complexity of alternative splicing and low circRNA abundance, current computational methods unfortunately fail to accurately detect BSJs while also producing exceedingly unsatisfying full-length assemblies. Therefore, the problem of *in silico* assembly of circRNAs, from highly sensitive BSJ identification to full-length circRNA reconstruction, remains largely unresolved.

Here we present TERRACE (accura**T**e ass**E**mbly of circ**R**NAs using b**R**idging and m**AC**hine l**E**arning), a new tool for assembling full-length circRNAs from paired-end total RNA-seq data. TERRACE stands out by assembling circRNAs accurately without relying on annotations, a feature absent in most existing tools. TERRACE starts with a fast, light-weight algorithm to identify back-spliced reads (i.e., reads containing BSJs). We realize that the key to assembling circRNAs is to correctly “bridge” the three fragments in a back-spliced read. We formulate this task as seeking paths in a weighted splice graph that can connect the three fragments and that its “bottleneck” weight is maximized, and solve this formulation using an efficient dynamic programming algorithm. TERRACE also features a new machine-learning model that is trained to assign confidence scores to assembled circRNAs which we show outperforms abundance-based ranking.

## 2 Methods

TERRACE takes the alignment of paired-end total RNA-seq reads and, optionally, a reference annotation as input. TERRACE first identifies back-spliced reads (Section 2.1), each of which will be assembled into a set of candidate, full-length circular paths in the underlying splice graph (Sections 2.2 and 2.3). The candidate paths, optionally augmented by the annotated transcripts, are subjected to a selection process followed by a merging procedure to produce the resultant circRNAs (Section 2.4). A score function is learned that can assign a confidence score to each assembled circRNA (Section 2.5). TERRACE is outlined in Fig. 1.

**Fig. 1:**
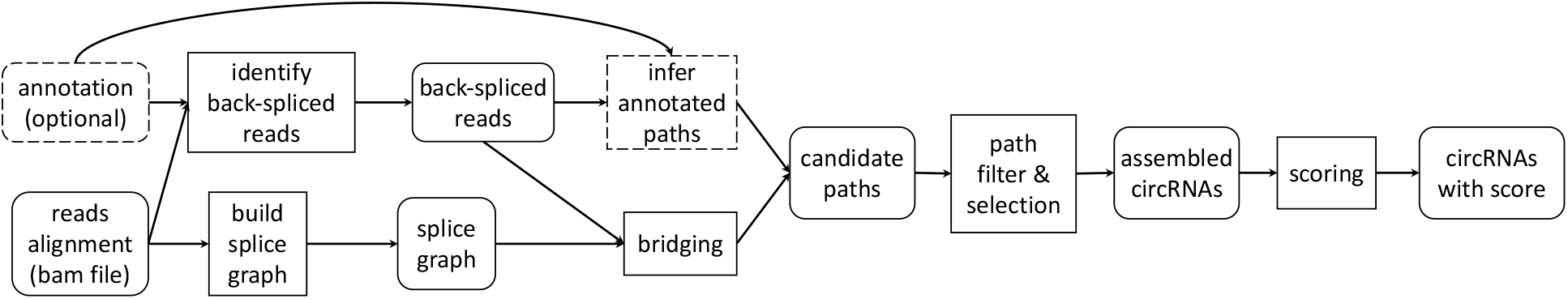
Outline of TERRACE. Rounded boxes represent data and data structures. Rectangles represent procedures. Dashed boxes indicate optional.

### 2.1 Identification of Back-Spliced Reads

The back-spliced reads (see Fig. 2) are pivotal in detecting circRNAs. TERRACE identifies back-spliced reads from two sources: *chimerically aligned* reads in the input alignment, and by a new, light-weight, junction-targeted mapping algorithm. A chimerically aligned read is a special class of reads where different portions of it are aligned to different locations of the reference genome. These reads are indicative of structural variation, including BSJs. In the bam format, one of its alignments is recorded as *primary* and others as *supplementary* alignments. TERRACE looks for chimerically aligned reads with only one supplementary alignment. Let *R1* and *R2* be the two ends of a paired-end read, where we assume *R1* is chimerically aligned, with its primary and supplementary alignment being denoted as *R1*.*primary* and *R1*.*supple* respectively. A pattern often exists in the CIGAR strings if *R*_1_ contains a BSJ. An example is given in Fig. 2, where the CIGAR strings of *R1*.*primary* and *R1*.*supple* are 30H70M and 30M70S, respectively. The 30 matched base pairs in the supplementary alignment complement the 30 unaligned portion (*hard clipped*, denoted by an ‘H’) in the primary alignment while the 70 matched base pairs in the primary alignment complement the 70 *soft clipped* portion in the supplementary alignment. Such a complementary relationship strongly indicates a BSJ. TERRACE collects chimerically aligned reads satisfying this relationship as back-spliced reads.

**Fig. 2:**
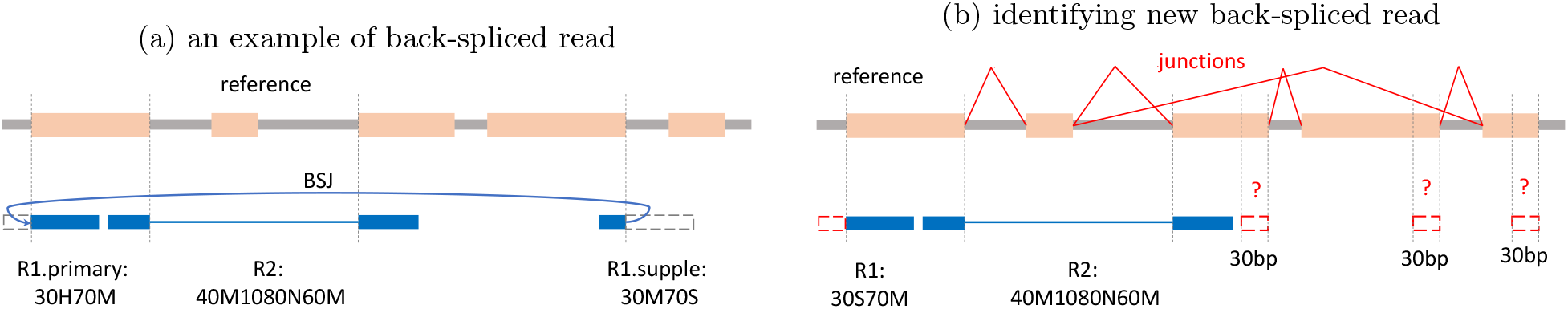
(a), the CIGAR strings of the 3 segments in a back-spliced read. (b), the soft clipped sequence (dashed box in red) will be compared with the reference sequence next to a splicing position of the same length.

TERRACE also implements a new method to identify more back-spliced reads whose chimeric property are not captured by the aligner. First, splicing positions are extracted from read junctions (represented by an ‘N’ in the CIGAR; a junction specifies two splicing positions). Additional splicing positions are also identified from the annotated transcripts given a reference transcriptome is provided. The collected splicing positions will be used as supporting evidence for BSJs. Next, the reads that have soft clips at either end greater than a threshold (a parameter of TERRACE with a default value of 15) are considered candidates for back-spliced reads. The sequence of the soft clipped region, denoted as *S*, will be remapped to the reference genome at a splicing position to identify a significant match. More specifically, for each candidate splicing position that may form a BSJ with the soft clipped region (depending on the relative locations of *R1* and *R2*), we extract a sequence of the same length as *S* from the reference genome, denoted as *T*, and calculate the Jaccard index of the two sets of kmers in *S* and *T*, where by default *k* = 10. If the Jaccard is greater than a threshold (0.9 by default) and there does not exist another such *T*, TERRACE will use it and create a supplementary alignment record for *S* by mapping it to *T*. Now we have a new back-spliced read that can be treated as the same way as those identified from chimerically aligned reads.

### 2.2 Transforming Assembly to Bridging

A back-spliced read is presumably expressed from a circRNA; we aim for assembling its original circRNA for each back-spliced read. Recall that a back-spliced read *R* with ends *R1* and *R2* consists of three segments *R1*.*primary, R2*, and *R1*.*supple*, assuming *R1* contains the BSJ. We already know that these three fragments must be part of the original circRNA, and that the circRNA uses the BSJ to form its circular structure. What we still miss is how the three segments are connected in the circRNA, as the three segments may be aligned to distant portions of the reference genome. We refer to the task of closing the two gaps among the three segments in a back-spliced read as “bridging”: assembling the full-length circRNA now becomes bridging. See Fig. 3(b).

**Fig. 3:**
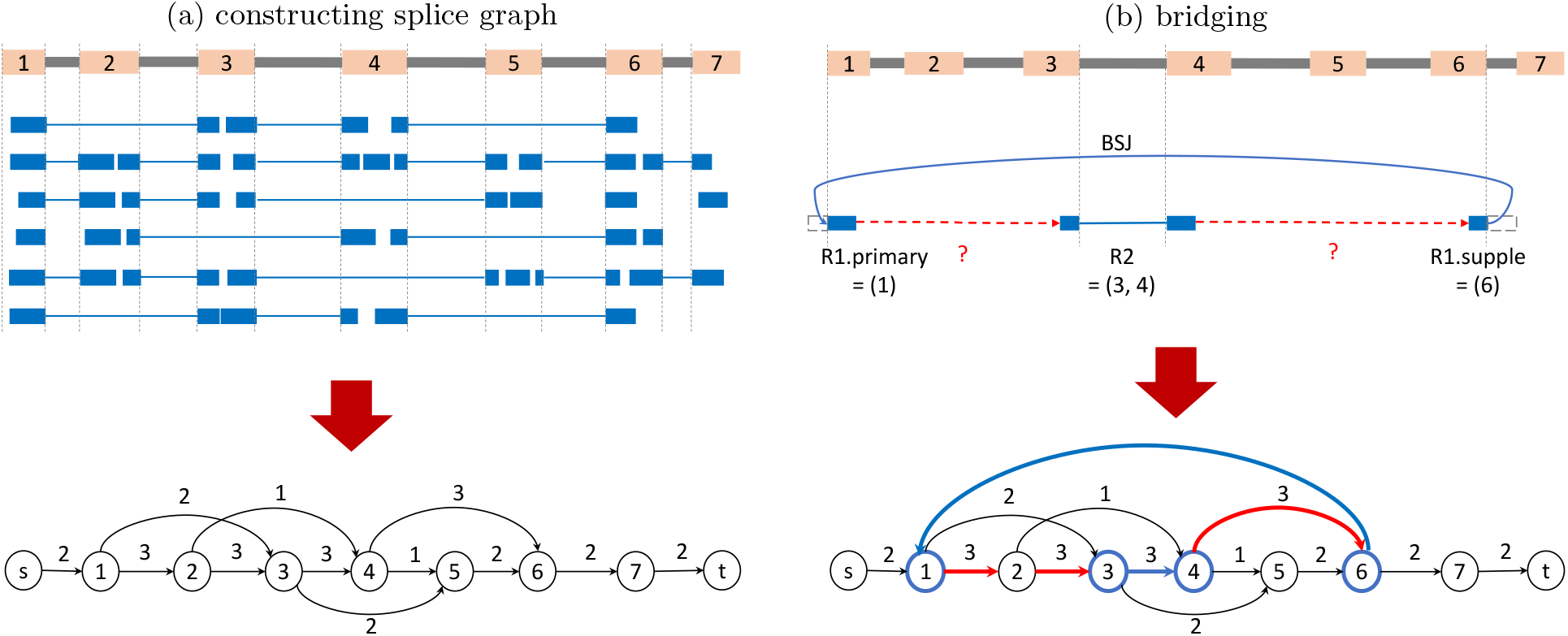
(a), constructing splice graph from reads alignment. (b), the 3 fragments namely *R1*.*primary, R2*, and *R1*.*supple* of a back-spliced read are represented as paths in the splice graph, which are (1), (3,4), and (6), respectively. The two optimal bridging paths, which maximize the “bottleneck” weight, are marked in red. The resulting full-length circular path for this back-spliced read is 1 ⟶ 2 ⟶ 3 ⟶ 4 ⟶ 6 ⟶ 1.

To formalize the bridging task, we introduce the underlying data structure: splice graph. See Fig. 3(a). Splice graph has been instrumental in studying alternative splicing and in assembling (linear) transcripts. It is a weighted directed graph, denoted as *G* = (*V, E, w*), that organizes the splicing and coverage information in the read alignment of a gene locus. To construct *G*, junctions from reads and annotated transcripts (if provided) are collected and their splicing positions will be used to partition the reference genome into (partial) exons and introns, in which the partial exons will be the vertices *V* of *G*. A directed edge *e* ∈ *E* is placed between vertex *u* and vertex *v* if there exists a read that spans *u* and *v*; the weight *w*(*e*) of edge *e* will be the number of such reads. A source vertex *s* is added and connected to any vertex *u* with in-degree of 0 using weight *w*(*s, u*) = ∑_*v*:(*u,v*)∈*E*_ *w*(*u, v*); a sink vertex *t* is also added and any vertex *v* with out-degree of 0 will be connected to *t* with weight *w*(*v, t*) = ∑_*u*:(*u,v*)∈*E*_ *w*(*u, v*).

The three fragments of a back-spliced read can be represented as three paths in the splice graph *G*. The bridging task now involves finding two *bridging paths* in *G* that connect the three known paths. Solving this results in a single path that threads the 3 fragments, which forms the circRNA together with the BSJ. Due to alternative splicing and sequencing/alignment errors, multiple possible bridging paths exist. We therefore need a characterization for better bridging paths and an efficient algorithm to calculate them.

### 2.3 Formulation and Algorithm for Bridging

Let *A* = (*a*_1_, *a*_2_, …, *a*_*i*_), *B* = (*b*_1_, *b*_2_, …, *b*_*j*_), and *C* = (*c*_1_, *c*_2_, …, *c*_*k*_) be the 3 paths in *G* corresponding to the 3 fragments of a back-spliced read. We aim to find the “best” paths in *G* from *a*_*i*_ to *b*_1_ and from *b*_*j*_ to *c*_1_. We adopt a definition we proposed in reconstructing the entire fragment of paired-end RNA-seq reads [35,15]. The idea was to seek a path whose “bottleneck weight” is maximized, which is effective for selecting the path with the strongest support and for excluding false paths due to errors which often contain edges with a small weight. Formally, we define the *score* of a path *p* as the smallest weight over all edges in *p*. The formulation is to find a path *p*_1_ from *a*_*i*_ to *b*_1_ and a path *p*_2_ from *b*_*j*_ to *c*_1_ such that the score of *p*_1_ and that of *p*_2_ are maximized. The optimal *p*_1_ and *p*_2_ can be calculated independently (we assume paths *A, B*, and *C* are vertex-disjoint; otherwise, they can be either bridged trivially or bridging is not possible in which case this read will be discarded). An efficient dynamic programming algorithm can be designed to find optimal *p*_1_ and *p*_2_. Please refer to [35] for details. The programming algorithm can be extended to produce suboptimal paths as candidates (by default TERRACE calculates top 10 optimal paths for each back-spliced read), which will be combined with additional information for selection.

### 2.4 Selection of Candidate Paths

Let *P* be the set of candidate full-length circular paths for a back-spliced read. We apply some heuristic procedures to filter false-positive paths. If a path *p* ∈ *P* contains a vertex (partial exon) in which a region of length larger than a threshold (by default 10 base pairs) is not covered by any read then *p* will be removed from *P*. For every pair of paths *p, q* ∈ *P*, if an intron of *p* is fully covered by an exon of *q*, then *q* will be removed. This procedure helps to filter out paths with anticipated intron retentions. If there exists a path *p* ∈ *P* with bottleneck weight higher than a chosen threshold *c* (by default *c* = 1), all paths in *P* with bottleneck weight smaller than *c* are discarded. This procedure aims to remove less reliable paths when more reliable ones exist. We use *P*_1_ to denote the set of survived paths.

If a reference transcriptome is provided, TERRACE will then identify more candidate full-length circular paths from it. See Fig. 4. We define an annotated (linear) transcript *q* is *compatible* with a back-spliced read *R* if both the BSJ and the splicing positions in the three fragments of *R* match the splicing positions of *q*. Unique compatible paths bounded by the BSJ will be collected as another set *P*_2_ of candidate full-length circular paths for *R*. If *P*_1_ ∩ *P*_2_*≠* ∅, the path in *P*_1_ ∩ *P*_2_ with maximized bottleneck weight will be picked; if *P*_1_ ∩ *P*_2_ = ∅ and *P*_1_*≠* ∅, we pick the path in *P*_1_ with maximized bottleneck weight regardless of *P*_2_ (i.e., we give higher priority to paths inferred from the read alignment rather than from reference annotation); otherwise, we will pick one path from *P*_2_ randomly if *P*_2_*≠* ∅. This selection procedure assigns one path to read *R* if *P*_1_ ∪*P*_2_*≠* ∅; otherwise *R* will be discarded. The selected full-length circular path is then transformed to a fully annotated circRNA by borrowing genomic coordinates from the reference genome.

**Fig. 4:**
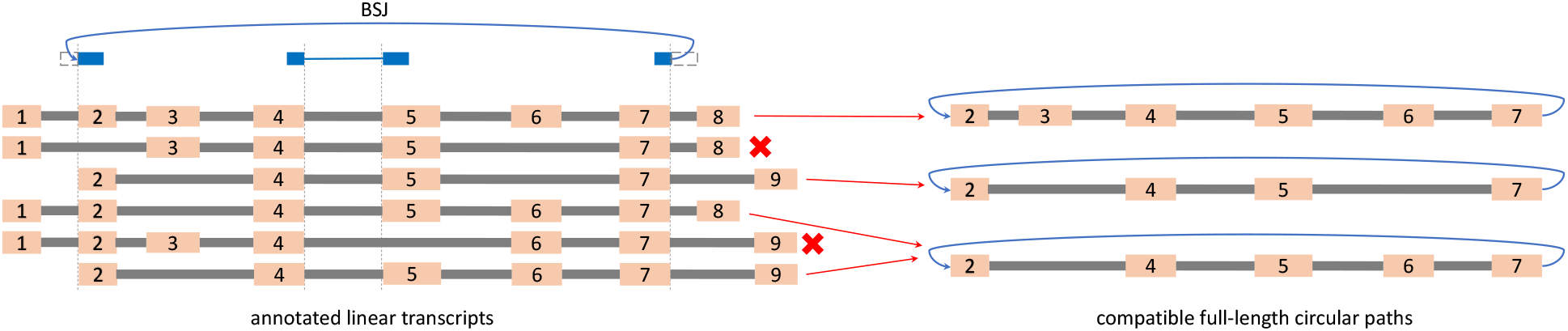
Identifying compatible full-length circular paths for a back-spliced read from annotated transcripts.

Note that multiple back-spliced reads can produce identical circRNA. We merge identical circRNAs to a single instance and the number of back-spliced reads generating this circRNA is recorded as its *abundance*. We observe that circRNAs may differ by a few base pairs at their BSJs but share the same intron chain. In such case, the circRNA with higher abundance is retained.

### 2.5 Scoring Assembled Circular RNAs

Assigning a confidence score to assembled circRNAs is desirable to ensure that those with higher scores are more likely to be true. Traditionally, abundance has served this purpose in transcript assembly as it shows a high correlation to the correctness of the assembled transcripts. We investigate whether a machine-learning approach could yield a more accurate scoring function. To this end, we extract 13 features to characterize each assembled circRNA, ranging from its abundance to the (average) length of the soft clips of back-spliced reads. Please refer to Appendix A for a detailed description of all features.

We use a random forest model trained on one tissue sample (brain) and tested it on other samples (see Section 3.1). We aim to train a model that is capable of generalizing across different tissues. This is challenging due to the variability between samples and the limited number of instances available. To fortify its stability, we incorporate the number of reference transcripts present in each instance (gene locus) as features, and use the assembled circRNAs by TERRACE with and without reference annotations to train the model. This approach is proven beneficial in generalizing, especially when the test set is markedly different from the training set or when the dataset is small (such as skeletal muscle). Additionally, this enables a single model to be applicable on circRNAs assembled both with and without annotations (instead of training two models).

## 3 Results

### 3.1 Experimental Setup

#### TERRACE

We implemented the method explained in Section 2 as a new circRNA assembler named TERRACE. TERRACE takes the alignment (in BAM format; can be produced by any RNA-seq aligner such as STAR [4] or HISAT [12]) of total RNA-seq reads as input and produces a list of full-length, annotated circRNAs in the GTF format. TERRACE is designed to assemble reliable circRNAs without the need for an annotation but comes with the flexibility of an optional reference annotation file.

#### Methods for Comparison

Although there are an abundance of methods that predict BSJs, very few of them produce the full-length annotation (please refer to CYCLeR paper [22] for a classification). Full-length circRNA assemblers that do not require a reference annotation are even rarer. We only identify CIRI-full as such a tool and it serves as the state-of-the-art in the field. We include CIRCexplorer2, CIRI-full and CircAST for comparison when the reference annotation is provided. CircAST needs to be provided with a list of BSJs from CIRI2 or CIRCexplorer2; we choose to use CIRI2 as the results are worse when CIRCexplorer2 is used. CYCLeR is not suitable for comparison since it necessitates both control total RNA-seq samples and circRNA-enriched samples, whereas our study exclusively focuses on total RNA libraries. Sailfish-cir and CIRCexplorer3 are tools that primarily target circRNA quantification and hence are not included.

#### Datasets

We use both real dataset and simulated dataset for evaluation. The real dataset is chosen from the isoCirc [30] paper which consists of 8 human tissue samples (lung, brain, skeletal muscle, heart, testis, liver, kidney, and prostate). Full-length circRNAs are cataloged for these samples using a combination of a reference gene annotation and long reads, which we use as ground-truth. The simulated dataset is generated using CIRI-simulator [5], previously used by several methods [6,34,11,20]. The expressed circRNAs are available in the simulation which we use as ground-truth. We simulate 10 samples and report the average performance.

#### Evaluation

We define an assembled circRNA to be *correct* if the coordinates of its BSJ and its intron-chain all exactly match a circRNA in the ground-truth. We then calculate *recall*, defined as the proportion of circRNAs in the ground truth that are correctly assembled, and *precision* which is the percentage of assembled circRNAs that are correct. It is common that a method exhibits high precision but low sensitivity, or vice versa. To still conclude in this case, we report *Fscore*, calculated as (2×*recall*×*precision*)*/*(*recall*+*precision*). We also draw the precision-recall curve for TERRACE. With the curve we can calculate an *adjusted precision* w.r.t. another method, defined as the precision of TERRACE at a point on the curve when its recall matches the recall of the compared method. We draw and compare two precision-recall curves for TERRACE, by using either the “abundance” or the “score” inferred by the random forest.

### 3.2 Comparison of Assembly Accuracy

Fig. 5 shows the accuracy of TERRACE and CIRI-full on the real dataset when the reference annotation is not provided. TERRACE yields a significantly higher number of correct circRNAs than CIRI-full across all samples (228% more on average) at a comparable precision (43% vs 40% on average). The Fscore of TERRACE is consistently higher than that of CIRI-full (22% vs 10% on average). Both the precision-recall curves of TERRACE are above CIRI-full, proving TERRACE’s much enhanced accuracy. Specifically, the average adjusted precision of TERRACE is 75% (using the random-forest curve; it is 72% using the coverage curve) comparing with CIRI-full at 40%.

**Fig. 5:**
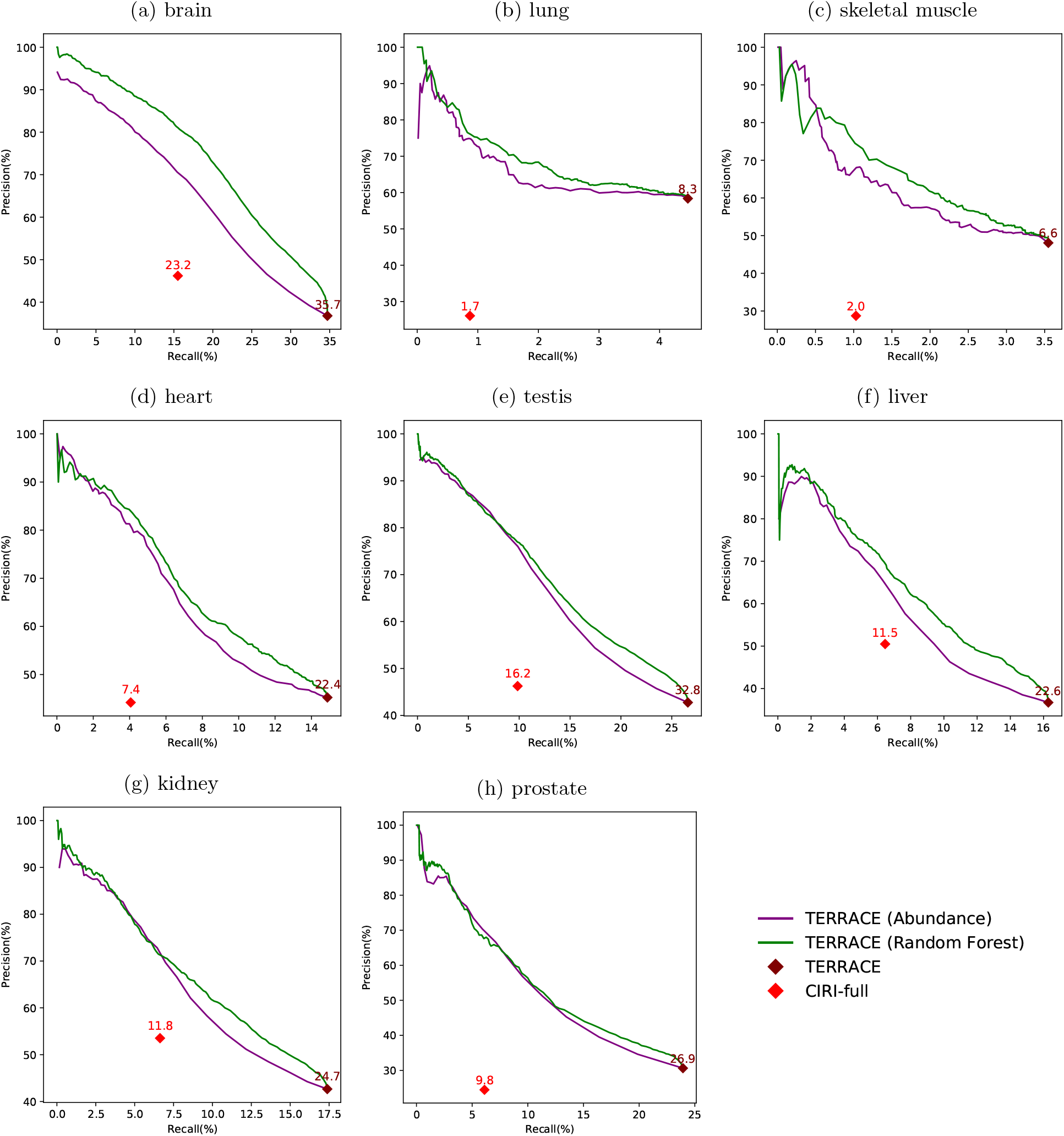
Comparison of assembly accuracy without annotation. Fscores (%) are indicated on top of data points.

The confidence scores produced by random forest result in a superior curve than that using abundance as the varying parameter. This improvement can be attributed to the machine learning model’s ability to learn a more effective and accurate scoring function from a broad class of features (including abundance). Of note, when the abundance is high, simply using it can identify correct circRNAs accurately in a performance similar to random forest (the low-recall regions in the figures). However, when the abundance becomes low, correct and incorrect circRNAs become indistinguishable by just using abundance. The trained scoring function effectively tackles this problem, ensuring a bigger improvement in precision in the high-recall regions.

Fig. 6 compares the accuracy of different methods running with an annotation. TERRACE reconstructs 18% more correct circRNAs than CIRCexplorer2 while obtaining higher precision on all samples. TERRACE assembles 272% more correct ones than CIRI-full at a comparable precision (42% vs 40% on average). CircAST exhibits higher precision, but at a cost of much reduced recall. Again, TERRACE obtains a higher Fscore than all other methods on all samples. Also, the precision-recall curves of TERRACE consistently lie well above the all other data points, demonstrating its improved accuracy. In particular, the average adjusted precision of TERRACE (using the random-forest curve) is 77% compared to CIRI-full at 40%, 50% compared to CIRCexplorer2 at 40%, and 86% compared to CircAST at 59%.

**Fig. 6:**
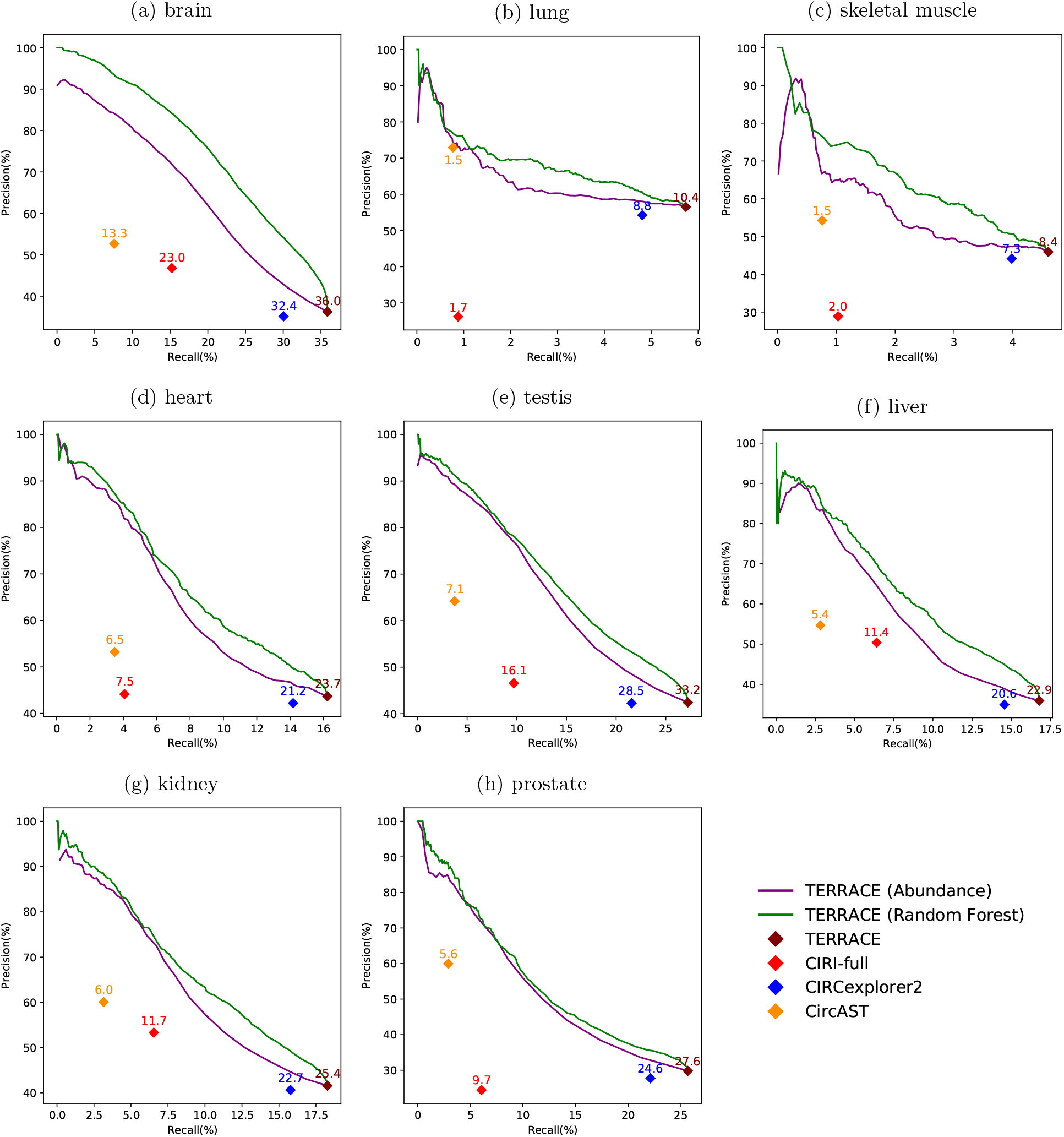
Comparison of assembly accuracy with annotation. Fscores (%) are indicated on top of data points.

Fig. 7 shows the accuracy of TERRACE compared to other tools on simulated data. When annotation is not provided, TERRACE identifies an average of 37% more correct circular transcripts and achieves better precision than CIRI-full. In the presence of annotations, TERRACE consistently outperforms CIRI-full and CIRCexplorer2 on all measures. TERRACE exhibits an average precision comparable to CircAST while maintaining much better recall, resulting in a much higher overall Fscore.

**Fig. 7:**
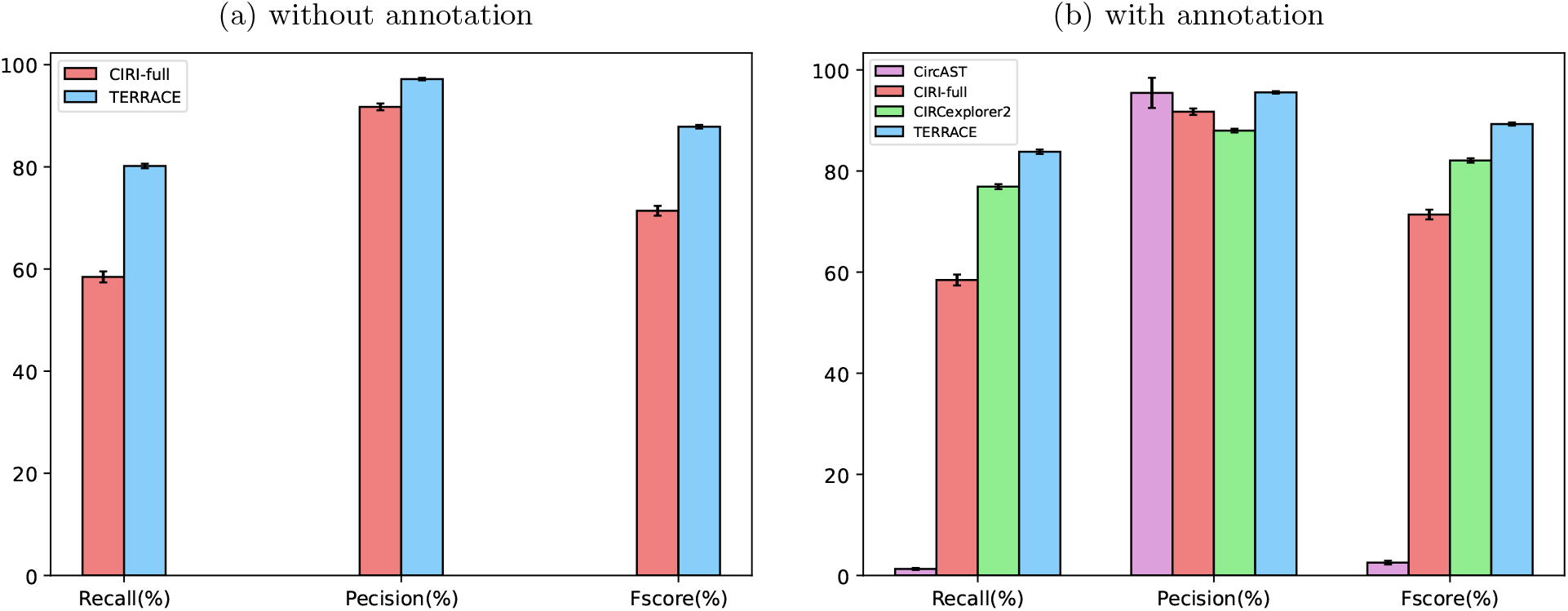
Average accuracy of different tools on simulated data. The error bars show the standard deviation over 10 simulated samples.

We realize that the recall rates of various methods including TERRACE are quite low on real data (Figs. 5 and 6), particularly in lung and muscle tissues. This trend is also evident in the precision values. One plausible explanation for this discrepancy could be linked to the use of annotated circRNAs from long reads data as the ground truth for evaluation. Upon analyzing the count of annotated circRNAs across the samples and comparing them to the limited number detected by various methods (refer to Appendix B), we hypothesize that long reads and short-reads total RNA-seq data (which is used here) capture divergent sets of expressed circRNAs. This may result in accurate predictions from short-reads being misclassified as incorrect, leading to an underestimation of both precision and sensitivity. As an evidence, we notice that the read coverage across the gene loci in muscle and lung samples is highly non-uniform: many reads cluster densely within a few number of gene loci, leaving other genes with sparse coverage. This causes all methods, including TERRACE, to construct much less number of correct circRNAs compared to the ground truth. The number of circRNAs annotated using long reads is in the similar range with other samples, so the coverage non-uniformity seems not to exist in the long reads dataset. Nevertheless, we emphasize that although the ground truth may not fully capture the absolute accuracy, it still serves as a fair benchmark for comparing the relative accuracy of different methods. Given the underestimated recall rates of TERRACE on the biological samples, we resort to simulations to illustrate that achieving high recall is possible in an unbiased setting. The performance of TERRACE on simulated samples strongly reinforces this assertion.

### 3.3 Comparison of Runtime and Memory Usage

Table 1 shows the CPU time (user time plus kernel time) of various tools on the real dataset. CIRCexplorer2 has the fastest execution time which is expected given it utilizes a more compact annotation file than the raw version. TERRACE is the second fastest, surpassing both CircAST and CIRI-full by a large margin. Overall, TERRACE delivers vastly superior accuracy within a reasonable processing time.

**Table 1:** CPU time (in minutes) for different tools on various datasets.

| sample | methods w/o annotation |  | methods with annotation |  |  |  |
| --- | --- | --- | --- | --- | --- | --- |
|  | CIRI-full | TERRACE | CIRCexplorer2 | CircAST | CIRI-full | TERRACE |
| lung | 265 | 31 | 0.14 | 249 | 266 | 28 |
| brain | 471 | 46 | 0.21 | 1234 | 489 | 49 |
| skeletal | 334 | 42 | 0.10 | 260 | 337 | 52 |
| heart | 380 | 35 | 0.10 | 488 | 389 | 38 |
| testis | 427 | 42 | 0.14 | 922 | 472 | 43 |
| liver | 330 | 32 | 0.08 | 423 | 345 | 33 |
| kidney | 352 | 33 | 0.10 | 554 | 370 | 35 |
| prostate | 253 | 54 | 0.24 | 229 | 307 | 61 |
| average | 352 | 39 | 0.14 | 545 | 372 | 42 |

Table 2 shows the peak memory usage of various tools on the real dataset. While TERRACE may not excel in peak memory usage, it still operates within an acceptable range and outperforms CIRI-full significantly. It is important to highlight that the both running time and peak memory usage values of TERRACE are very similar regardless of whether an annotation is provided (i.e., the use of annotation does not significantly impact its computational efficiency).

**Table 2:** Peak memory usage (in GB) for different tools on various datasets.

| sample | methods w/o annotation |  | methods with annotation |  |  |  |
| --- | --- | --- | --- | --- | --- | --- |
|  | CIRI-full | TERRACE | CIRCexplorer2 | CircAST | CIRI-full | TERRACE |
| lung | 10.1 | 16.9 | 0.14 | 0.07 | 11.0 | 16.7 |
| brain | 80.4 | 11.2 | 0.16 | 0.11 | 81.1 | 11.4 |
| skeletal | 22.2 | 19.8 | 0.14 | 0.10 | 22.8 | 20.2 |
| heart | 30.8 | 11.0 | 0.14 | 0.61 | 31.4 | 11.1 |
| testis | 74.8 | 5.4 | 0.15 | 0.09 | 76.2 | 5.6 |
| liver | 271.9 | 17.7 | 0.14 | 0.08 | 31.7 | 17.9 |
| kidney | 32.0 | 10.3 | 0.14 | 0.08 | 33.1 | 10.1 |
| prostate | 21.7 | 23.4 | 0.15 | 0.08 | 22.4 | 23.6 |
| average | 68.0 | 14.5 | 0.15 | 0.15 | 38.7 | 14.6 |

## 4 Conclusion and Discussion

The substantial growth in research dedicated to circRNAs in recent years underscores their significance in biology and medicine. Despite the abundance of experimental and computational techniques designed for circRNA detection, inherent limitations persist within these methods. Current experimental protocols often require special enrichment of libraries for accurate detection while computational methods are hindered by their dependence on annotation and inability to reconstruct full-length molecules. TERRACE made a significant advancement towards closing this critical gap. TERRACE utilized a fast algorithm that effectively detects previously overlooked back-splicing junctions. A bridging system, which we originally proposed for improving (linear) transcript assembly, was re-designed to reconstruct the full-length circRNAs. Instead of using abundance for ranking, TERRACE learned a better score function with a broader class of informative features. TERRACE outperforms existing tools drastically, especially in scenarios where annotations are unavailable. We anticipate widespread adoption of TERRACE, particularly in studies involving species lacking well-annotated transcriptomes.

Further improvements can be made for TERRACE. The precision-recall curves (in Figs. 5 and 6) are not satisfactory. We would investigate the extraction of more features and advanced learning approaches. For instance, we can incorporate the count of splicing positions or partial exons supporting a BSJ as extra features. Additionally, we could explore training a model with complete sequences of *bona fide* circRNAs, possibly sourced from established circular RNA databases, to better differentiate between accurate and erroneous ones. An LSTM model may be used for such sequence-based training. We also consider extending TERRACE to incorporate additional types of data. Leveraging long reads and/or circRNA-enriched libraries for detection may reveal less obvious circRNAs, enhancing the assembly accuracy.

## Availability

The source code of TERRACE is freely available at https://github.com/Shao-Group/TERRACE. The scripts, evaluation pipelines, and instructions that can be followed to reproduce the experimental results of this work is available at https://github.com/Shao-Group/TERRACE-test. The raw sequencing (fastq) files of the 8 samples is available at BIGD (accession number: PRJCA000751). The alignment files (by STAR) of these samples and the raw sequences and alignment files of the simulated data have been hosted at Penn State Data Commons (DOI: https://doi.org/10.26208/AZ99-RQ38).

## Acknowledgment

This work is supported by the US National Science Foundation (2019797 and 2145171 to M.S.) and by the US National Institutes of Health (R01HG011065 to M.S.). We thank Qimin Zhang for helpful input. We thank Fangqing Zhao and Jinyang Zhang for their assistance with CIRI-full.

### Appendix A

We elaborate the features used to train the scoring function described in Section 2.5 and provide our insights on why they can be informative. The features are characterizing an assembled circRNA *x*.

1. Coverage. This is the number of back-spliced reads that produce *x*. Intuitively, a circRNA supported by a higher number of back-spliced reads is more likely to be correct.
2. Count of additional back-spliced reads that produce *x*. By additional, we mean those back-spliced reads identified by TERRACE but missed by the aligner. The argument for including this as a feature is similar to the intuition in 1, i.e., a higher abundance of reads is an evidence for real circRNAs.
3. Sum of soft clip lengths of the back-spliced reads that produce *x*. Back-spliced read with a longer soft clip is more likely to be correctly aligned to the correct junction. Hence, circular RNAs characterized by longer soft clips are more likely to be genuine.
4. Sum of bridging path scores of reads that produce *x*. Recall that the score of a path represents its bottleneck weight. Paths with higher scores receive support from more reads, and may be an important factor in distinguishing between true and false instances.
5. Sum of the count of full-length candidate paths from back-spliced reads that produce *x*. By default, TERRACE considers the top 10 bridging paths and selects a set of them using some filtering criteria (see Section 2.4) to be further analyzed. The greater the number of paths in this selected set, the more likely it is to deviate from choosing the correct path.
6. Count of bridging path type of back-spliced reads producing *x*. The path type refers to whether the selected path is inferred from read alignments or reference transcripts and if the path length is within the range of insert size. When selecting a bridging path, we normally would want to assign a higher priority to paths inferred from read alignment than from the reference annotation, aiming to construct novel circRNAs. However, if the set of paths inferred from reads is empty, sometimes a path from the annotation (if provided) may help recover a correct circRNA specially when the coverage within a region is low. These features may provide guidance to the machine learning model to make accurate decisions.
7. Number of exons in *x*. In certain high-coverage areas of specific samples, numerous false circRNAs with a large number of exons are detected due to the presence of many small splicing sites. Hence, this information can be an important factor.
8. Total length of exons in *x*. We observe that some circRNAs have extended exon lengths, which are misleading and primarily caused by intron retentions. Therefore, considering the total exon length as a feature could prove valuable in making decisions.
9. Maximum length of exons in *x*. Intuition similar to 8.
10. Minimum length of exons in *x*. Intuition similar to 8.
11. Total number of reads in the region where *x* is identified. We observe a few instances where a false circRNA is supported by many reads, primarily in some high coverage regions of certain samples. Involving the number of reads (both ordinary and back-spliced) as a feature may help to normalize this bias.
12. Total number of reference transcripts in the region where *x* is identified. The splicing positions from reference transcripts (when annotation provided) influences the identification of additional chimeric reads and adds to the set of paths to be considered for bridging. Therefore, the number of annotated transcripts in a region may serve as a useful feature for learning a better score.

### Appendix B

**Table 3:**
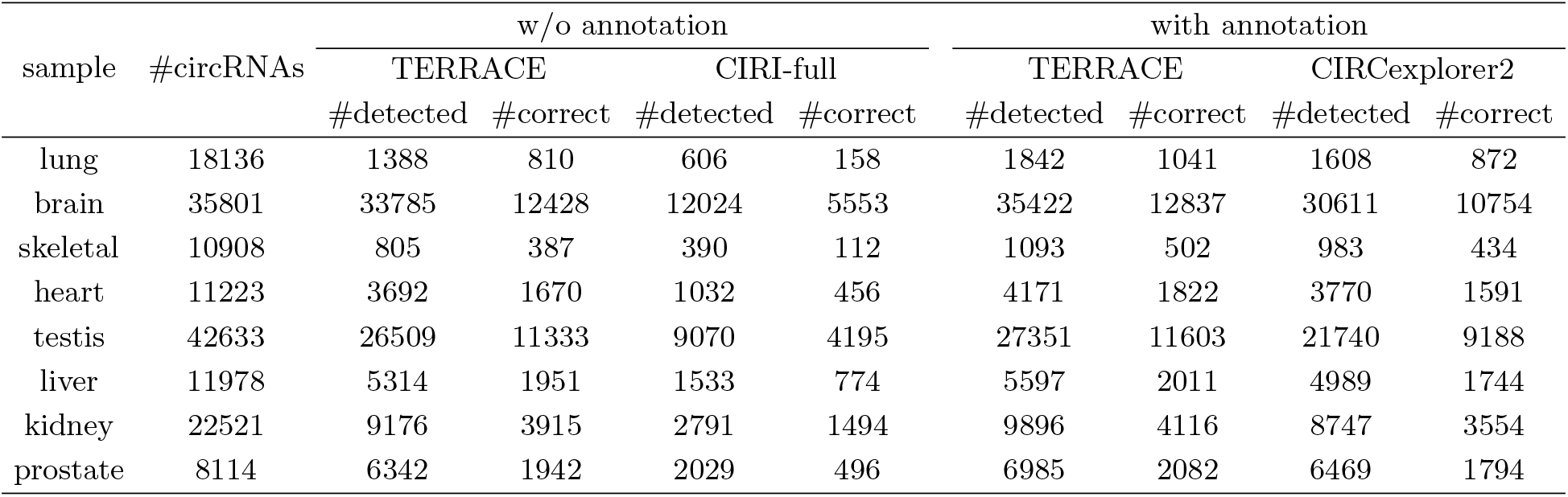
Number of circRNAs produced from long-reads in the isoCirc paper (which we use as ground truth for evaluation), number of circRNAs assembled and correctly identified by TERRACE, CIRI-full, and CIRCexplorer2.

